# Design of PEG-based hydrogels as soft ionic conductors

**DOI:** 10.1101/2024.06.17.599239

**Authors:** Gabriel J. Rodriguez-Rivera, Fei Xu, Madeline Laude, Vani Shah, Abbey Nkansah, Derek Bashe, Ziyang Lan, Malgorzata Chwatko, Elizabeth Cosgriff-Hernandez

## Abstract

Conductive hydrogels have gained interest in biomedical applications and soft electronics. To tackle the challenge of ionic hydrogels falling short of desired mechanical properties in previous studies, our investigation aimed to understand the pivotal structural factors that impact the conductivity and mechanical behavior of polyethylene glycol (PEG)-based hydrogels with ionic conductivity. Polyether urethane diacrylamide (PEUDAm), a functionalized long-chain macromer based on PEG, was used to synthesize hydrogels with ionic conductivity conferred by incorporating ions into the liquid phase of hydrogel. The impact of salt concentration, water content, temperature, and gel formation on both mechanical properties and conductivity was characterized to establish parameters for tuning hydrogel properties. To further expand the range of conductivity available in these ionic hydrogels, 2-acrylamido-2-methyl-1-propanesulfonic acid (AMPS) was incorporated as a single copolymer network or double network configuration. As expected, conductivity in these ionic gels was primarily driven by ion diffusivity and charge density, which was dependent on hydrogel network formation and swelling. Copolymer network structure had minimal effect on the conductivity which was primarily driven by counter-ion equilibrium; however, the mechanical properties and equilibrium swelling was strongly dependent on network structure. The structure-property relationships elucidated here enables the rationale design of this new double network hydrogel to achieve target properties for a broad range of applications.

## INTRODUCTION

Conductive hydrogels are increasingly used in many applications such as soft electronic devices, actuators, wearable sensors, touch panels, batteries, energy storage devices, biomedicine, and sustained drug release systems [1-5], which can be categorized into two main modes including electronic and ionic conductivity. Electronic conductivity relies on electron transport through metal or unsaturated bonds in molecules such as polyaniline, and polypyrrole.[6, 7] Although highly conductive, incorporation of electronically conductive particles or copolymers can limit applications due to issues with biocompatibility, gel heterogeneity, or processing.[8, 9] Ionic conductivity, in turn, relies on the movement and solubility of ions incorporated into the aqueous phase of the hydrogel. Although ionic hydrogels typically have lower conductivity than electrically conductive materials [10-13], the conductivity still exceeds that of relevant biological tissues.

In an ionic hydrogel, the conductivity is impacted by the polymer network and the liquid composition. First, the ion concentration and types of ions present in the network are known to impact conductivity. Zhou et al. incorporated additional ion molecules in the hydrogel network to enhance both conductivity and mechanical properties.[5] Incorporating microstructure that contains a high concentration of ions such as in the case of hydroxypropyl cellulose aggregates has also been shown to impact conductivity.[5] For biomedical applications, ion selection is limited due to the ions present in the extracellular fluid and physiological concentrations of ions. Given the free exchange of ions between the hydrogel and the surrounding extracellular fluid, we would expect that the ion concentration and identity will typically equilibrate after implantation in the body, resulting a limit on the range of ionic conductivity of neutral hydrogel polymer backbones. To shift this equilibrium to increase the hydrogel conductivity, ionic polymers such as polyelectrolytes can be incorporated into the networks and increase the conductivity due to the presence of counterions. A variety of polyelectrolyte hydrogels have been studied to elucidate the impact of polycation and/or polyanion species. Lee et al. compared ten different chemically distinct gels including zwitterionic, cationic, anionic, and nonionic gels, and found that the presence of counterions in cationic and anionic hydrogels led to higher conductivity in low electrolyte concentration solutions than that in zwitterionic hydrogels [14]. Beyond conductivity, the mechanical properties of the hydrogel are critical constraints in the design of medcial devices and components. Significant effort has been made to improve the mechanical properties of conductive hydrogels to achieve these requirements, such as tuning gel chemistry to introducing additional bonding modalities (e.g., hydrogen bonding, metal ion coordination.[15, 16] Another successful strategy is the incorporation of a second network or double network that can increase mechanical strength and toughness due to interpenetrating network entanglement and efficient energy dissipation.[17, 18] Despite recent advances, it continues to be challenging to achieve both high conductivity and desired mechanical properties.

To expand the utility of hydrogels as soft ionic conductors, this work aims to elucidate key hydrogel structure-property relationships to enable the rational design of hydrogels with target conductivity and mechanical properties. A polyethylene glycol (PEG)-based hydrogel was selected given its established biocompatibility, soft tissue-like mechanical properties, and broad utility in biomedical applications.[19-21] In our previous study, we synthesized polyether urethane diacrylamide (PEUDAm), a more hydrolytically stable PEG macromer than poly(ethylene glycol) diacrylate (PEGDA).[22, 23] Similar to PEGDA, PEUDAm is a neutral polymer; therefore, its conductivity depends solely on the ions in the aqueous phase. To expand the range of conductivity, 2-acrylamido-2-methyl-1-propanesulfonic acid (AMPS) was selected as an anionic comonomer due to its prior use in biomedical, agriculture, and food science applications.[24] Previous studies evaluated the effect of crosslinker concentration in PEG-AMPS copolymers which simultaneously impacts swelling and conductivity.[25] In this study, we decouple those effects by fabricating hydrogels with single networks, copolymer networks, and interpenetrating networks (IPN), respectively (**Figure 1**). The effect of ion concentration, water content, network structure, and gel formation on both conductivity and mechanical properties were studied in this report. The resulting structure-property relationships provide a polymer toolbox for researchers to enhance ionic conductivity while also maintaining adequate mechanical properties of the hydrogel.

**Figure 1.**
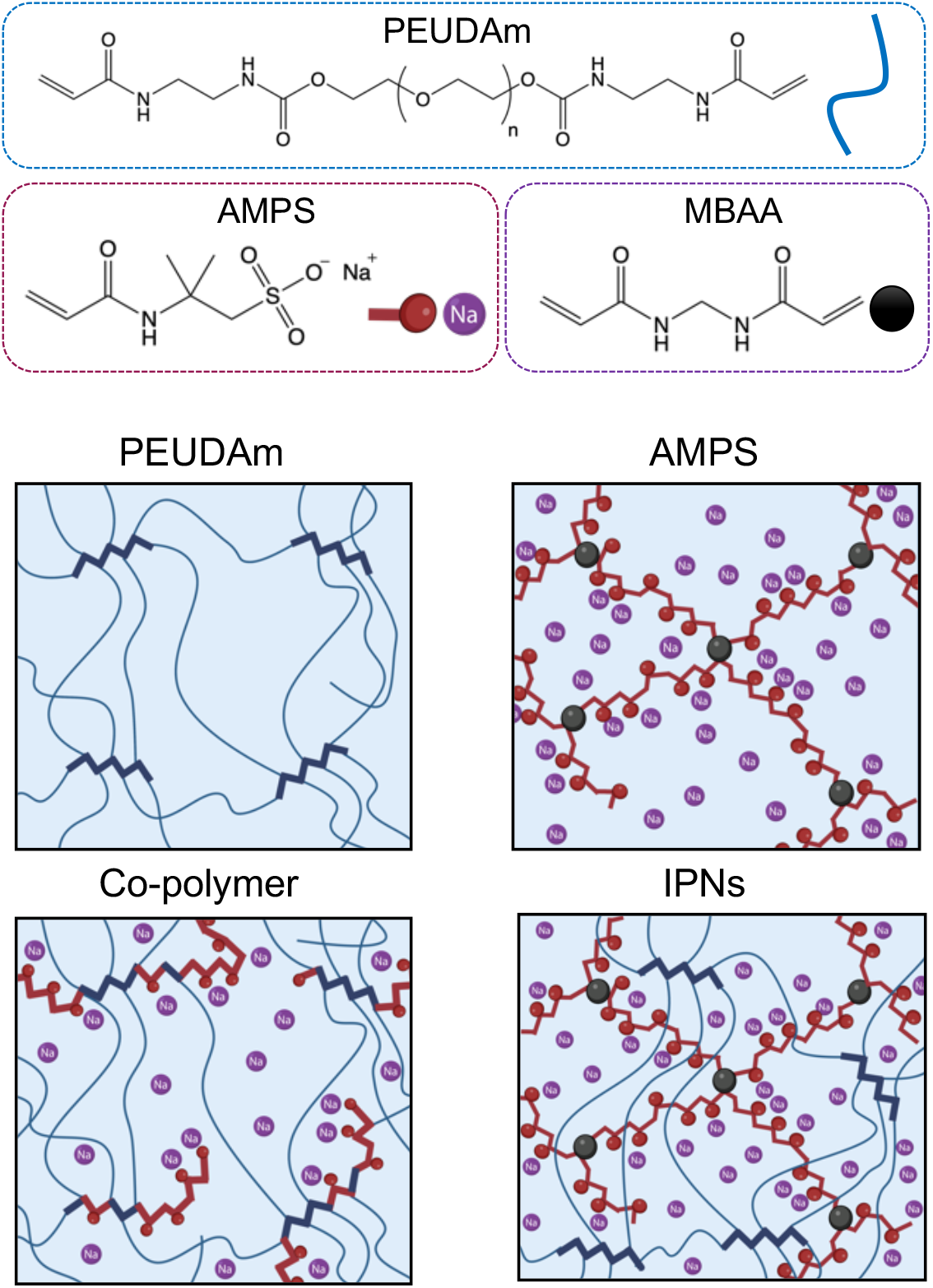
Chemical structure of macromer and monomers used in ionic hydrogels with schematic of single network, copolymer network and interpenetrating network (IPN) structures.

## RESULTS AND DISCUSSION

### Impact of ions concentration and gelation on conductivity of single-network hydrogels

First, the impact of gelation and ion concentration on conductivity of single-network hydrogels was studied. It was noted that the water content at swelling equilibrium between AMPS and PEUDAm hydrogels were significantly different (19.3 ± 0.7 for PEUDAm and 23.4 ± 1.1 for AMPS in DI water, 16.4 ± 0.3 for PEUDAm and 12.0 ± 1.0 for AMPS in 0.9% saline, Supporting Information **Table S2**). This difference was attributed to differences in crosslink density and the shifting balance of osmotic pressure with ionic strength of the medium and backbone.[26] To minimize this effect, a fixed water content was used to study the effect of macromer chemistry and gelation on conductivity. Macromer solutions and hydrogels were tested at a weight ratio of 12:1 (water : dried polymer, Q∼92 wt%). As shown in **Figure 2**, both hydrogels and solutions exhibited a proportional increase of conductivity with increasing NaCl concentration from 0 to 1.35 wt%. In contrast to the intrinsic conductivity of AMPS hydrogel, the conductivity of the neutral PEUDAm hydrogel exclusively depended on the NaCl concentration. (**Figure 2A-B)** The conductivity of the PEUDAm hydrogel precursor solution was similar to the PEUDAm hydrogel at low salt concentrations from 0 to 0.45 wt%; with only a small difference observed at higher concentrations of 0.9 wt% and 1.35 wt% (**Figure 2C)**. This indicates that the hydrogel network had minimal effects on ion diffusivity, which is the primary mode of ionic conductivity in these hydrogels.

**Figure 2.**
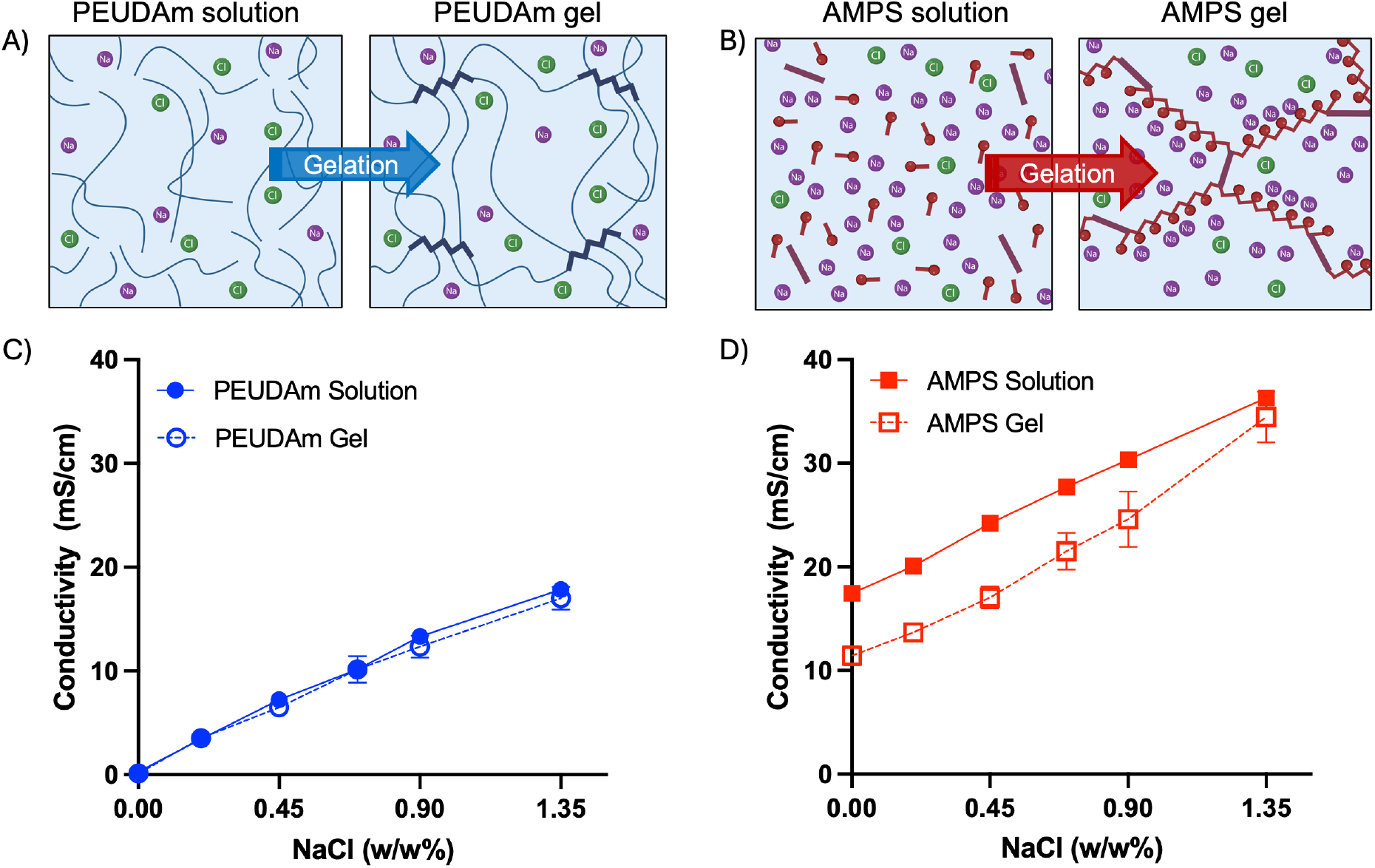
Conductivity as a function of ion concentration and gelation. A) Schematic of impact of gelation on conductivity of neutral PEUDAm hydrogel and B) charged AMPS hydrogel. C) Impact of NaCl concentration and gel formation on conductivity of neutral PEUDAm solution and hydrogel, and D) AMPS solution and hydrogel. Mean and standard deviation are presented (n = 3). Error bars shorter than the size of the symbols are not displayed.

In contrast, AMPS hydrogels displayed a higher initial conductivity in DI water (11 mS/cm) due to its charged backbone. As expected, increasing NaCl concentration in the swelling medium also resulted in a linear increase in conductivity (**Figure 2D**). There was a marked decrease in conductivity of the AMPS hydrogel as compared to its hydrogel precursor solution at different salt concentrations. This difference was attributed to a decrease in molecule mobility of the ionic monomer and Manning condensation in hydrogel network at salt concentrations up to 0.9 wt%. However, the AMPS hydrogel and precursor solution displayed similar conductivity at the highest salt concentration of 1.35%. This may indicate that NaCl ions became the main contributors to hydrogel conductivity (**Figure 2D**). In both cases, a two-way ANOVA was used to confirm the statistically significant differences as salt content increases, between the solutions and hydrogels and the interaction between both factors.

### Effect of swelling on hydrogel conductivity

The impact of swelling on hydrogel conductivity was determined by assessing single-network hydrogels with controlled water content.(**Figure 3**) Hydrogels were dried prior to the addition of a calculated amount of DI water or saline (0.9 wt%). For the neutral PEUDAm hydrogel, only saline was used to hydrate the dried gels with different weight ratios given that the conductivity in DI water was minimal. With increased swelling in saline, the conductivity of PEUDAm hydrogels increased from 4.9 ± 0.5 mS/cm to 12.6 ± 0.6 mS/cm (**Figure 3C**). This increase in conductivity was attributed to the increase in ion mobility as the hydrogel network expands. (**Figure 3A**) For AMPS hydrogels, DI water and saline were used to hydrate dried hydrogels, respectively. The starting ionic conductivity of AMPS hydrogels in both DI water and saline at a swelling ratio of 3 was similar (24.1 ± 0.7 mS/cm in DI water and 24.6 ± 0.5 mS/cm in saline), due to the presence of counter ions in the polyelectrolyte hydrogels. (**Figure 3D**) The conductivity of AMPS hydrogels decreased with increased swelling in both DI water and saline, but the effect was markedly less in saline as compared to DI water. This decrease was attributed to the reduced charge density at higher water content as the charged backbone expands with swelling. This effect is mitigated when swelling with saline due to the ion concentration in the swelling medium. (**Figure 3B**).

**Figure 3.**
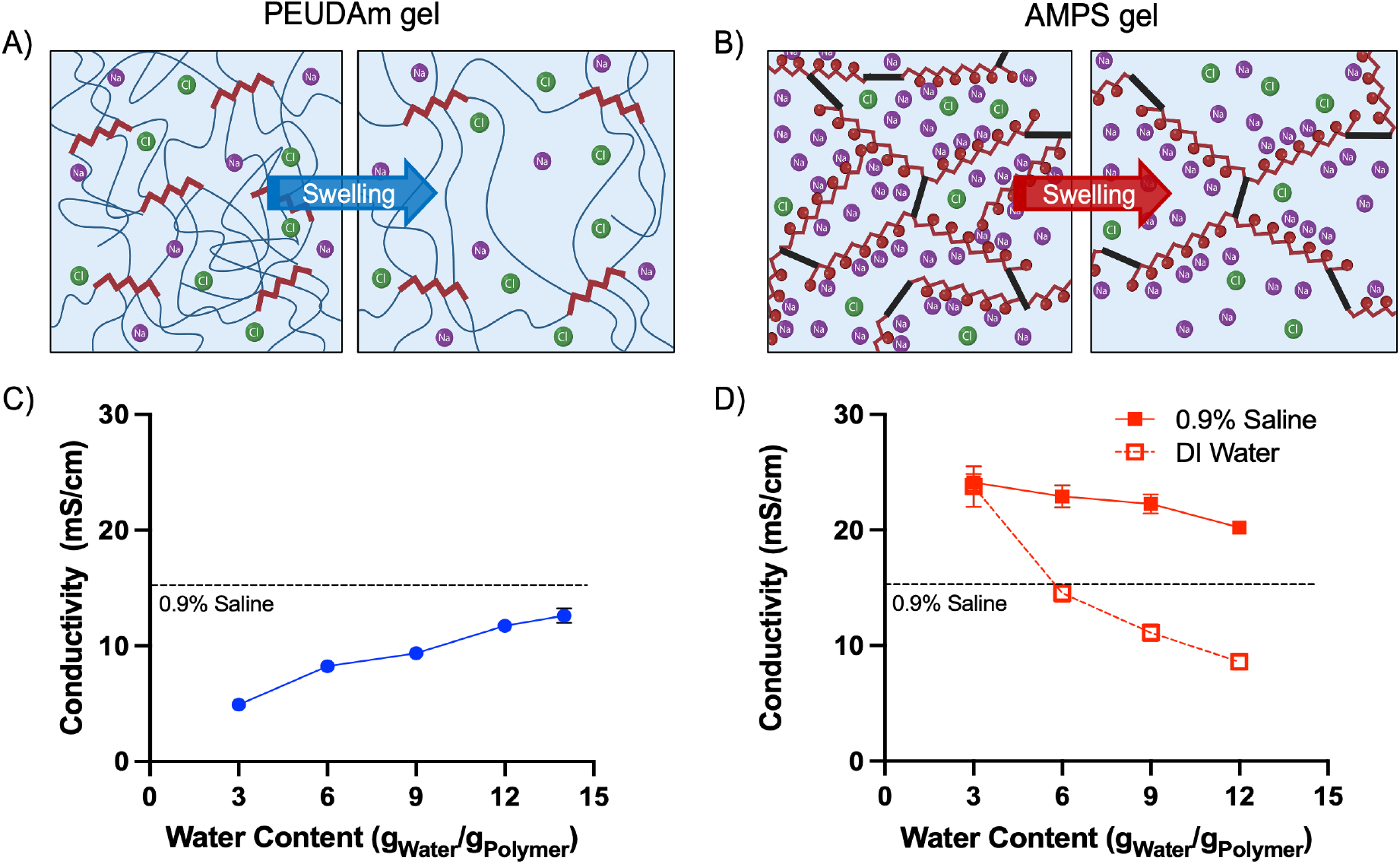
Conductivity as a function of hydrogel swelling. A) Schematic of impact of swelling on conductivity of PEUDAm hydrogel and B) AMPS hydrogel. C) Conductivity of PEUDAm hydrogel as a function of saline solution (0.9% NaCl) added to the dry polymer. D) Conductivity of AMPS hydrogel as a function of the solution (saline, DI water) added to the dry polymer. Mean and standard deviation are presented (n = 3). Error bars smaller than the size of the symbols are not shown.

### Effect of copolymer network design on conductivity

Although AMPS hydrogels displayed marked increases in conductivity, the hydrogel was brittle which limits its use in several applications. Indeed, tensile testing could not be completed because the AMPS hydrogel specimens either slipped from the grips or broke at the grips at less than 10% strain. To combine the conductivity of AMPS hydrogel with the mechanical property ranges available with PEUDAm, we explored two copolymer network designs: copolymerization or interpenetrating networks (IPNs). Co-polymer and IPN hydrogels were fabricated at equal mass composition (50:50 PEUDAm:AMPS). The conductivity of PEUDAm hydrogels increased similarly with the addition of the AMPS as a co-monomer or second network. It was also notable that no significant differences (two-way ANOVA analysis) between the copolymer network and IPNs hydrogels under the same water content and saline concentrations (**Figure 4**), indicating similarity of the main conduction mechanism caused by the diffusion of free ions in solution. In addition, at similar mass composition in both copolymer and IPN networks, the molar ratio of AMPS to PEUDAm is 87 to 1, resulting in most likely block AMPS structures in the kinetic chain that would be comparable to the IPN kinetic chain. This would indicate that either approach could be used to improve hydrogel conductivity. The preliminary data showed an increase in the conductivity of the PEUDAm hydrogel when combined with AMPS using either a copolymer or an IPN (**Figure 4**).

**Figure 4.**
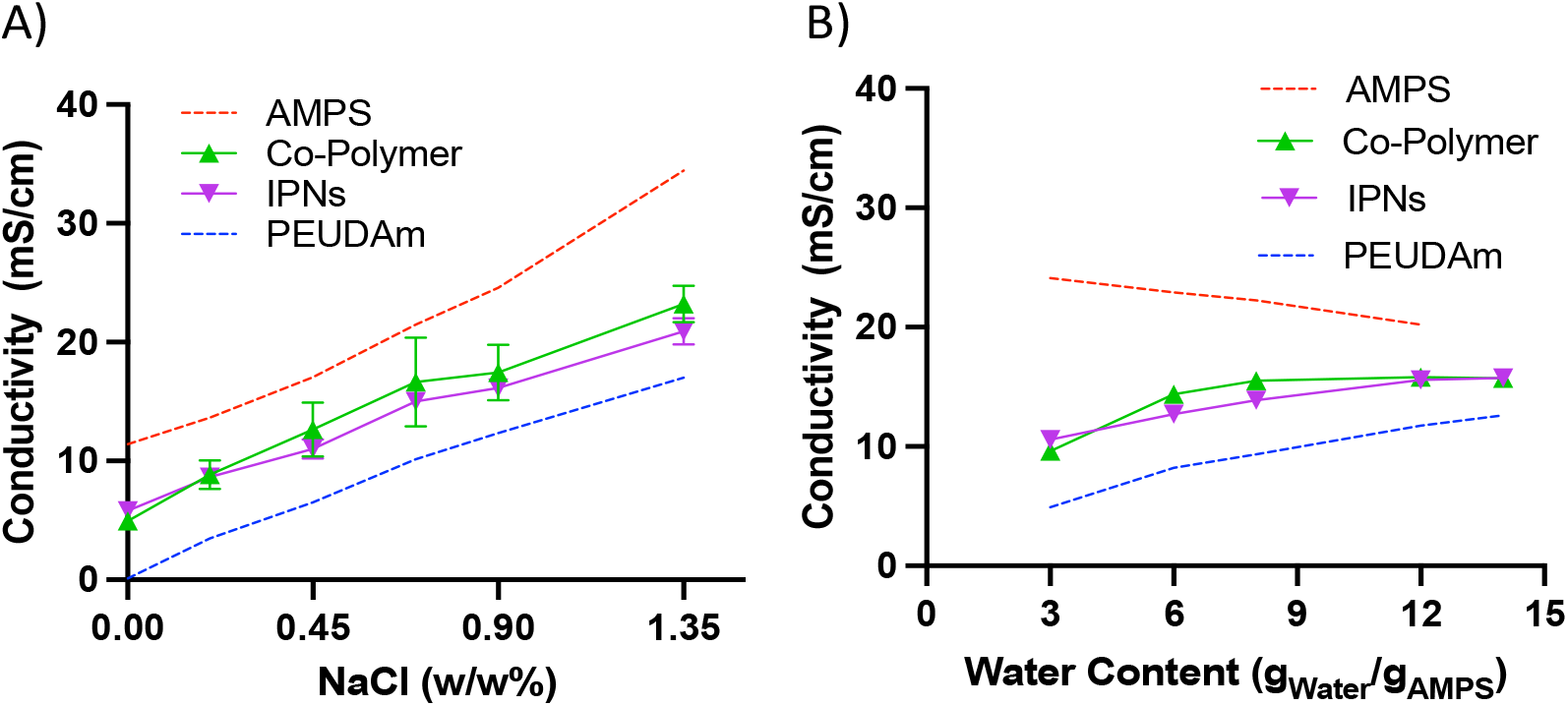
Effect of copolymer network structure on hydrogel conductivity. A) Conductivity of single network, copolymer, IPN hydrogels as a function of NaCl concentration at fixed water content. The water content of the hydrogels was 12 grams of solution per gram of dried polymer (water : polymer = 12 : 1). B) Conductivity of hydrogel compositions as a function of water content at fixed NaCl concentration (0.9% NaCl). Mean and standard deviation are presented (n=3). Error bars shorter than the size of the symbols are not displayed.

### Hydrogel properties at equilibrium swelling

All hydrogels displayed similar gel fractions (> 90%). The equilibrium swelling ratio of AMPS hydrogels was less than PEUDAm at similar polymer concentration (20 w/w%), **Figure 5A**. However, the AMPS showed a significant increase in swelling when equilibrated in DI water. This is in agreement with previous reports comparing different polyelectrolyte hydrogels, including PEGMA and AMPS (11). The main reason for the increase in swelling in water is the repulsion of the negative charge on the backbone, which is also observed on other anionic and cationic monomers (11). The increase in NaCl ions reduces the electrostatic repulsion of the anionic hydrogels resulting in gel shrinkage. When the AMPS monomers were copolymerized with PEUDAm, the effect was an equilibrium swelling ratio that was intermediate to the AMPS and PEUDAm single network hydrogels. When the AMPS was incorporated as a second network with PEUDAm, a small increase in swelling was observed as compared to the PEUDAm (**Figure 5A**). This may indicate that the crosslink density of the AMPS network was lower than in the single network hydrogel. As a result of the higher swelling, the conductivity of the IPN in equilibrium is lower than the copolymer hydrogel but still significantly higher than PEUDAm. (**Figure 5B**) As a result of the AMPS content and intermediate swelling, both copolymer and IPN hydrogels improved the conductivity of the PEUDAm, but not as high as the single network AMPS.

**Figure 5.**
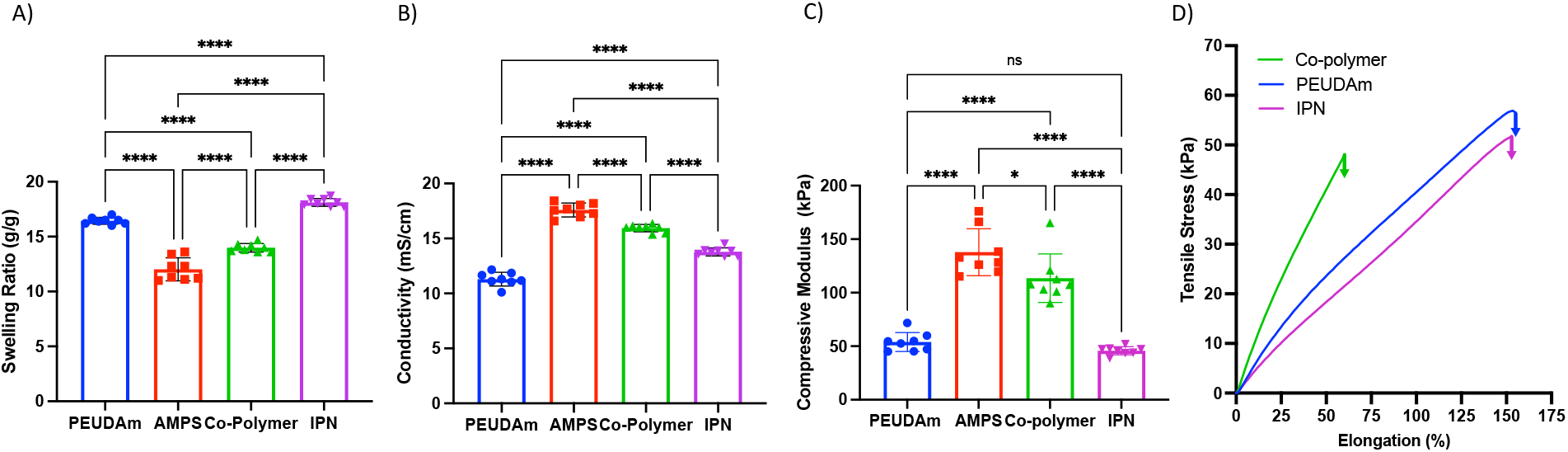
Hydrogel properties at equilibrium swelling in saline. A) Equilibrium swelling in saline (0.9% NaCl), n=8. B) Conductivity of hydrogels after swelling to equilibrium in saline, n=8. C) Compressive modulus of hydrogels after swelling to equilibrium in saline (1Hz, 1% strain), n=8. D) Representative stress vs. strain plot of hydrogels under uniaxial strain, n=3. Individual values, mean and standard deviation are presented (ns = not significant; ^*^ = p < 0.05; ^****^ = p < 0.0001)

### Impact of network structure on mechanical properties

One of the main findings of this work is that similar conductivity independent of network design can be achieved with similar composition and water contents (**Figure 4**). However, these hydrogels exhibit different mechanical properties at equilibrium (**Figure 5C-D**). On the other hand, PEUDAm single networks achieved elongations above 150% and modulus within the 50 kPa range. As a copolymer network, the resulting hydrogel exhibited an increase in stiffness and reduction in ultimate elongation. However, when the AMPS is added as an independent, second network, there was minimal impact on the hydrogel stiffness and elongation, indicating that the PEUDAm hydrogel retains most of its ability to dissipate stress in a more cooperative way with the AMPS network. It should be noted that the trends observed with the AMPS will likely translate to other polyelectrolyte monomers.

### Effect of temperature on hydrogel conductivity

The previous conductivity measures were completed at room temperature, ranging from 17 to 20°C. Since a physiological temperature of 37°C is necessary for utility in many biomedical applications, the effect of temperature (20-45°C) on conductivity was tested. There was a linear increase in conductivity with increased temperature (**Figure 6A**). The conductivity also followed an Arrhenius relationship within the studied range, (**Figure 6B**). This is also consistent with other PEG polyelectrolytes at different salts and polymer concentrations.[27] The PEUDAm hydrogel swollen in saline achieved a conductivity of 18.7 mS/cm, which is 3 times higher than highly conductive tissues (e.g.; myocardium ∼ 6 mS/cm) at the same temperature (**Figure 6A**). The single and double network hydrogels containing AMPS exhibited even higher conductivities at physiological temperatures.

**Figure 6.**
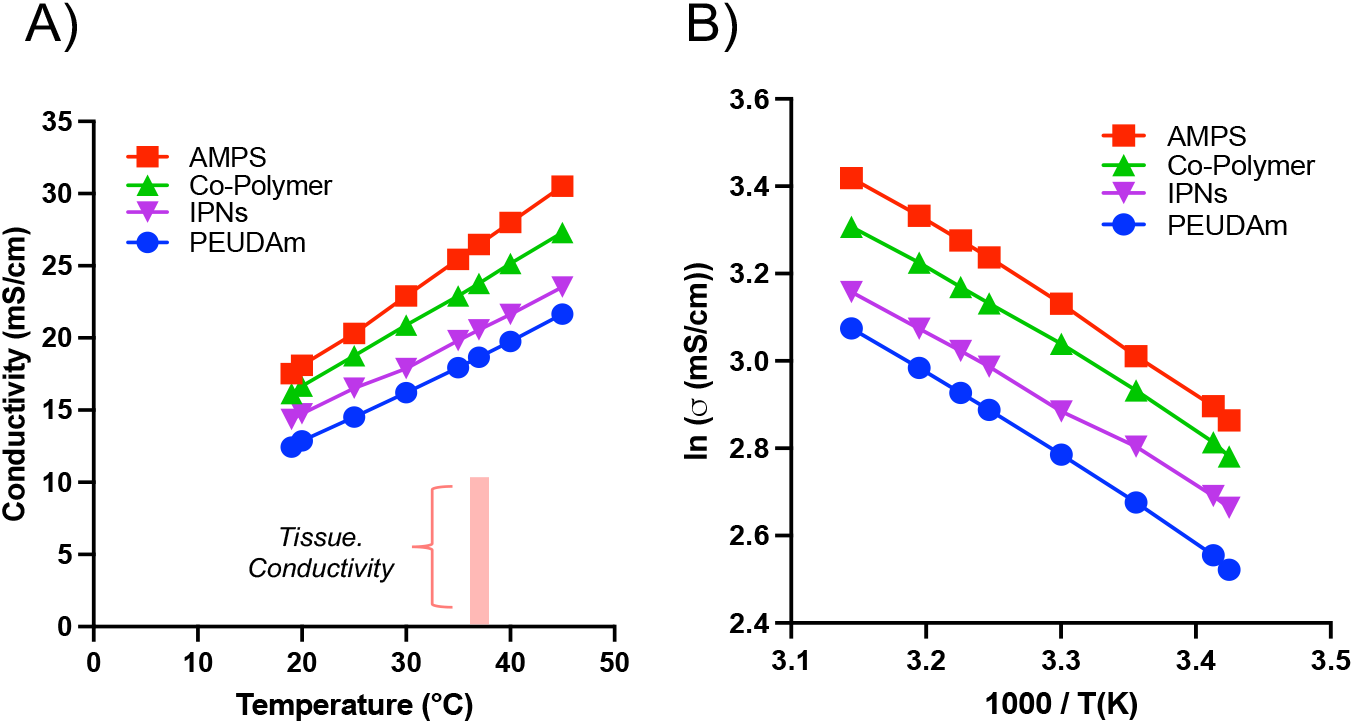
Effect of temperature on hydrogel conductivity. A) Conductivity as a function of temperature. The hydrogel ionic conductivity at physiological temperature (37°C) is higher than conductive tissues. B) The conductivity of the hydrogel follows an Arrhenius-type relation between room temperature and physiological temperatures.

## CONCLUSIONS

In this paper, we utilized a series of hydrogels to elucidate key structure-properties relationships that can be used to design PEG-based hydrogels with target ionic conductivity and mechanical properties. We isolated individual effects of gelation, ion concentration, water content, temperature, and network chemistry/structure on hydrogel conductivity. Gelation had minimal effect on the conductivity of the neutral PEUDAm hydrogel indicating that the swollen hydrogel network did not limit ion diffusivity. In contrast, a decrease in conductivity was observed upon gelation of the AMPS polyelectrolyte networks in accordance with other reports that describe ion condensation effects. As expected, a proportional increase in conductivity was observed in both PEUDAm and AMPS single-network hydrogels with increased ion concentration in the swelling medium. Conductivity also increased with increased swelling of PEUDAm hydrogels in saline due to an increase in ion diffusivity; whereas, swelling in the polyelectrolyte AMPS hydrogel resulted in a decrease in conductivity due to a decrease in charge density. Under similar swelling and ion concentrations, co-network and IPNs hydrogels showed similar conductivity, indicating that the concentrations of charge groups are the main driver in conductivity rather than network structure. However, copolymer structure affected mechanical properties and equilibrium swelling of the hydrogels. Elucidation of these key structure-property relations enable the rational design of conductive hydrogels with target conductivity, swelling, and mechanical properties for a broad range of biomedical applications.

## EXPERIMENTAL

### Materials

Poly(ethylene glycol) (PEG, Sigma Aldrich, No. 81275, Mn = 20000 g/mol), carbodiimidizole (CDI, Alfa Aeser No. A14688), anhydrous dichloromethane (DCM, from Sigma No. 34856), ethylene diamine (EDA, Sigma Aldrich), triethylamine (Sigma Aldrich), acryloyl chloride (Sigma Aldrich), 2-acrylamido-2-methyl-1-propanesulfonic acid (AMPS, Sigma Aldrich, 99%), triethylamine (Sigma Aldrich, > 99.5%), N,N-methylene-bis-acrylamide (MBAA, Sigma Aldrich), potassium carbonate (K_2_CO_3_, Sigma Aldrich), ether (Sigma Aldrich), Irgacure 2595 (Sigma Aldrich), sodium chloride (NaCl, Sigma Aldrich) were all used as received. Distilled deionized (DI) water were used for all experiments. The conductivity of DI water and NaCl solutions used for experiments was included in Supporting Information **Table S1**.

### PEUDAm Synthesis

PEUDAm was synthesized using a three-step process as reported previously.[23] In the first functionalization, PEG (20 kDa, 1 equiv., 1.74 mmol) was dissolved in DCM (300 mL) and reacted with CDI (15 equiv., 26.1 mmol) under nitrogen purge for 2 hours then allowed to react under stirring for 24 hours. The reaction was then quenched with water and the solvent was removed to yield PEG-CDI product. PEG-CDI (1 equiv.) was then reacted with EDA (15 equiv.) for 24 hours to yield the amine capped PEG macromer, PEG-EDA. Triethylamine was then added to PEG-EDA (1 equiv. in 250 mL DCM) with acryloyl chloride (4 equiv.) and allowed to react under stirring for 48 hours. The reaction was then quenched with K_2_CO_3_and the product precipitated in 1000 mL ice-cold ether. The final PEUDAm structure was confirmed with proton nuclear magnetic resonance (H^1^ NMR) spectra collected on a 400 MHz Bruker NMR. H^1^ NMR (CDCl_3_): δ = 6.94 (broad s, 2H, -C-N*H*-CH_2_-), 6.28 (dd, 2H, *H*_2_C=CH-C-), 6.17 (m, 2H, H_2_C=C*H*-C-), 5.85 (broad s, 2H, -C-N*H*-CH_2_-), 5.61 (dd, 2H, *H*_2_C=CH-C-), 4.20 (m, 4H, -H_2_C-C*H*_*2*_-O-), 3.65 (m, 1720H, -O-C*H*_*2*_-CH_2_-), 3.34 (m, 4H, -CH_2_-C*H*_*2*_-NH-).

### Hydrogel fabrication and characterization

PEUDAm was synthesized using a three-step process as reported previously and detailed in the supplemental information.[23] Single network hydrogels were fabricated with the PEUDAm (20 kDa, 20 wt%) macromer or AMPS monomer (20 wt%). For AMPS hydrogel networks, MBAA was used as a crosslinker (0.87 wt%). Bulk single network hydrogels were fabricated by dissolving PEUDAm or AMPS in 0.9% saline mixed with 0.1 wt% of the photoinitiator Irgacure 2595. For preparation of co-polymer network, PEUDAm (20 wt%) macromer was combined with AMPS monomer (20 wt%) in 0.9% saline mixed 0.1 wt% of the photoinitiator Irgacure 2595. Hydrogel precursor solutions were pipetted between glass plates with 1.5 mm spacers and UV cured on each side. PEUDAm hydrogels were cured for 6 min on each side; whereas, AMPS and copolymer hydrogels were cured for 30 mins on each side. For the IPN hydrogels, the single network of PEUDAm was fabricated as previously stated, followed by drying and swelling in AMPS monomer solution (8.33 wt%) with 0.1 wt% of Irgacure 2595 overnight prior to photocuring for 30 min on each side. A 12 ×volume of the AMPS solution to dry mass of the PEUDAm first network hydrogel yielded 1:1 mass ratio of AMPS and PEUDAm in the final IPN hydrogel.

To assess the equilibrium swelling ratio and gel fraction, 8 mm discs were punched from the hydrogel sheet immediately after curing and dried for 24 hours in a vacuum oven. Mass of dried hydrogels was measured (W_i_) and then the hydrogels were swollen to equilibrium in saline for 24 hours followed by swelling in DI water. Three washes with DI water were performed to remove the saline before equilibrium swelling in DI water. The swollen hydrogels were weighed (W_s_), then dried for 24 hours and reweighed (W_d_). The sol fraction was calculated as (W_i_-W_d_)/W_i_. The equilibrium volumetric swelling ratio, Q, was calculated from the equilibrium mass swelling ratio: W_s_/W_d_.

### Conductivity testing

The hydrogel impedance was determined using electrochemical impedance spectroscopy (MTZ-35, BioLogic). Briefly, the hydrogel discs were placed between two electrodes with an applied alternate current sweep from 1 to 10^6^ Hz. The minimum impedance was used to determine the resistivity. The resistance (R) of the hydrogel was determined when the impedance plateaued in the bode plot. Conductivity (σ) was calculated by Equation 1, where A is the electrode area (A) and L is the hydrogel thickness (L).

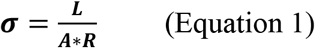

The conductivity of the hydrogel precursor solution was determined using a handheld conductivity probe (Oakton CON 450).

### Mechanical Testing

Hydrogels of each composition were fabricated as described above and tested at equilibrium swelling conditions in DI water. Bulk hydrogels were cut to dog-bone shapes using a custom 3D printed device as described by Nelson et al.[28] Testing was performed using a Dynamic Mechanical Analyzer (RSA III, TA Instruments). Hydrogel specimens (n=3) were strained to fracture at a rate of 0.1 mm/sec. Only the specimens that fractured in the gauge length and not at the grips were utilized for analysis. Specimen ultimate elongation, ultimate tensile strength, and modulus (between 5-15% strain) were calculated. For compressive modulus measurements, hydrogel disks (D = 8 mm, T = 1.5 mm) were punched from hydrogel sheets and swollen to equilibrium prior to testing. For compression testing, unconstrained specimens underwent a dynamic frequency sweep test at 1% strain using a dynamic mechanical analyzer (RSAIII, TA Instruments) equipped with a parallel-plate compression clamp. The linear viscoelastic range for each hydrogel formulation was determined using dynamic strain sweeps with an initial static force of 0.1 N. A specific strain value (typically ranging from 0.09 to 0.14%) was then chosen within the linear viscoelastic range and used for a sequential constant-strain frequency sweep test. The frequency sweep tests were run at frequencies ranging from 0.79 to 79 Hz, and the compressive storage modulus was calculated at 1 Hz. All mechanical tests were performed at room temperature.

### Effect of temperature on hydrogel conductivity

The previous conductivity measures were completed at room temperature, ranging from 17 to 20°C. Since a physiological temperature of 37°C is necessary for utility in many biomedical applications, the effect of temperature (20-45°C) on conductivity was tested. There was a linear increase in conductivity with increased temperature (**Figure S1A**). The conductivity also followed an Arrhenius relationship within the studied range, (**Figure S1B**). This is also consistent with other PEG polyelectrolytes at different salts and polymer concentrations.[27] The PEUDAm hydrogel swollen in saline achieved a conductivity of 18.7 mS/cm, which is 3 times higher than highly conductive tissues (e.g.; myocardium ∼ 6 mS/cm) at the same temperature (**Figure S1A**). The single and double network hydrogels containing AMPS exhibited even higher conductivities at physiological temperatures.

### Statistical Analysis

Quantitative characterization of hydrogel properties is expressed as mean ± standard deviation. Analysis of variance (ANOVA) and computations were performed using GraphPad Prism version 9.0.2 at the significance levels of p < 0.05. For graphs where the standard deviation is smaller than the bullet size, the error bars are not displayed by the software.

## ASSOCIATED CONTENT

### Supporting Information

Supporting information is available free of charge at http://pubs.acs.org/doi/xx.xxxx.

## ACKNOWLEDGMENTS

We thank Dr. Nate Lynd at The University of Texas at Austin for access to the electrochemical impedance equipment in his lab.

## AUTHOR INFORMATION

### Author contributions

- Conceptualization: GJRR, ECH, MC
- Methodology: GJRR, ECH
- Investigation: GJRR, ML, VS, AN, ZL
- Visualization: GJRR, ML
- Supervision: ECH
- Writing—original draft: GJRR, ECH
- Writing—review & editing: ECH, MC, FX, DB

