## Supporting Information for "Design of PEG-based hydrogels as soft ionic conductors"

**Table S1**. Conductivity of distilled deionized (DI) water and NaCl solution

| | DI water (μS/cm) | NaCl solution (mS/cm) | | | | | |
|---|---|---|---|---|---|---|---|
| | | 0.2% | 0.45% | 0.7% | 0.9% | 1.1% | 1.35% |
| Conductivity | 0.8 ± 0.1 | 4.0 ± 0.1 | 8.2 ± 0.1 | 13.0 ± 0.1 | 16.1 ± 0.1 | 19.4 ± 0.1 | 24.1 ± 0.1 |

*The conductivity measurement was conducted at 20.8 °C.

**Table S2**. Equilibrium swelling ratio (g/g) of hydrogels in DI water and 0.9% saline (n = 8)

| | PEUDAm | AMPS | Co-polymer | IPN |
|---|---|---|---|---|
| DI water | 19.3 ± 0.7 | 23.4 ± 1.1 | 21.1 ± 1.2 | 20.0 ± 0.4 |
| 0.9% saline | 16.4 ± 0.3 | 12.0 ± 1.0 | 14.0 ± 0.4 | 18.1 ± 0.4 |
